# Comparative Transcriptomic and Metabolomic Analyses in Response to Cold in Tartary Buckwheat (*Fagopyrum tataricum*)

**DOI:** 10.1101/278432

**Authors:** Jin Jeon, Jae Kwang Kim, Qi Wu, Sang Un Park

**Affiliations:** Department of Crop Science, Chungnam National University, 99 Daehak-ro, Yuseong-6 gu, Daejeon, 34134, Korea; Division of Life Sciences and Bio-Resource and Environmental Center, Incheon National University, Yeonsu-gu, Incheon 406-772, Korea; College of Life Science, Sichuan Agricultural University, 46 Xinkang Road, Ya’an, Sichuan 625014, China

**Keywords:** Anthocyanin, cold, metabolome, proanthocyanidin, tartary buckwheat, transcriptome

## Abstract

Plants recognize multiple environmental signals that lead to substantial changes in the regulation of primary and secondary metabolism in order to adapt to environmental stresses. In this study, we investigate the effects of cold on the metabolome and transcriptome of tartary buckwheat, focusing on the phenylpropanoid biosynthetic pathway. Using RNA-sequencing analysis of the cold-regulated transcriptome of buckwheat, we identified several phenylpropanoid biosynthetic transcripts that accumulated in response to cold. To confirm the transcriptome data, we analyzed the expression of the phenylpropanoid biosynthetic transcripts in cold-treated buckwheat and showed that all the phenylpropanoid biosynthetic transcripts were upregulated in cold-treated buckwheat seedlings with the exception of *FtDFR*. From the metabolic profiling based on the GC-TOF-MS analysis, we show that most of the sugars and their derivatives significantly increase in response to cold, while some of amino acids and their derivatives decrease in cold-treated plants. Some organic acids derived from the tricarboxylic acid (TCA) cycle increased in the cold-treated plants compared with in the plants grown at 25°C. In particular, the contents of anthocyanins and proanthocyanidins were significantly increased by cold treatment. In summary, these results indicate that the metabolome and transcriptome of tartary buckwheat are extensively affected by cold stresses.

**Highlight:** Using the RNA-sequencing data and the metabolic profiling analysis, we identified the changes that occur in the transcriptome and metabolome of tartary buckwheat in response to cold, focusing on the flavonoid biosynthetic pathway.

**Abbreviations:**

ANR: anthocyanidin reductase
ANS: Anthocyanin synthase
C4H: cinnamate 4-hydroxylase
CHI: chalcone isomerase
CHS: chalcone synthase
4CL: 4-coumarate:CoA ligase
DFR: dihydroflavonol reductase
DPPH: 2,2-diphenyl-1-picrylhydrazyl
F3H: flavanone 3-hydroxylase
F3’H: flavonoid 3’-hydroxylase
FLS: flavonol synthase
GC-TOFMS: gas chromatography-time of flight mass spectrometry
3GT: flavonoid 3-O-glucosyltransferase
HPLC: high-performance l iquid chromatography
LAR: leucoanthocyanidin
PA: proanthocyanidin
PAL: phenylalanine ammonia-lyase
RT: 3-O-rhamnosyltransferase
TCA: tricarboxylic acid.

## Introduction

Plants live in restricted spaces that are constantly exposed to various environmental stresses such as cold, drought, high salinity, high light/UV and heavy metals. During these stressful conditions, plants biosynthesize specialized metabolites to adapt to these environmental stresses. Many phenylpropanoid compounds are induced in plants by abiotic stresses (Dixon and Paiva, 1995). Wounding stimulates the accumulation of coumestrol, coumarin, chlorogenic acid and polyphenolic barriers such as lignin (Hahlbrock and Scheel, 1989; Bernards and Lewis, 1992). Wound-induced polyphenolic barriers may act directly as defense compounds. UV induces specific flavonol glycosides and sinapate esters in *Arabidopsis* and rutin and quercetin in buckwheat (Hectors *et al*., 2014; Huang *et al*., 2016). Anthocyanins and some flavonols accumulated in response to high light, cold and drought (Li and Cheng, 2013; Li *et al*., 2015; Nakabayashi *et al*., 2014).

Cold has a significant role in the regulation of primary and secondary metabolism in plants (Guy *et al*., 2008; Cook *et al*., 2004). Cook *et al*. (2004) used a GC-time of-flight MS metabolic profiling approach to identify which metabolome was altered in response to cold (Cook *et al*., 2004). Of the 434 metabolites monitored by this technique, 325 (73%) were found to increase in *Arabidopsis* Wassilewskija-2 (Ws-2) during cold conditions. The 114 metabolites (35%) including the amino acid proline and the sugars glucose, fructose, galactinol, raffinose and sucrose increased at least 5-fold in cold-treated plants. Cold affects the general phenylpropanoid and anthocyanin synthesis in apple skin, blood orange fruit, purple kale, buckwheat and maize seedlings by inducing the expression of flavonoid biosynthetic genes (Ubi *et al*., 2006; Crifo *et al*., 2012; Zhang *et al*., 2012; Li *et al*., 2015; Christie *et al*., 1994). The expression of five anthocyanin biosynthetic genes including *MdCHS, MdF3H, pDFR, MdANS* and *pUFGluT* was enhanced by cold treatment with the accumulation of anthocyanins in apple skin during ripening (Ubi *et al*., 2006). The anthocyanin levels in cold treated purple kale were approximately 50-fold higher than those in plants grown in a greenhouse (Zhang *et al*., 2012). The expression of the anthocyanin biosynthetic genes *BoC4H, BoF3H, BoDFR, BoANS* and *BoUFGT* and the transcription factor *BoPAP1* were enhanced by cold in purple kale. The anthocyanin content of blood orange fruit exposed to cold sharply increases more than 8-fold compared to that observed in the untreated samples (Crifo *et al*., 2012). The expression of anthocyanin biosynthetic genes (*CM1, PAL, CHS, DFR, ANS* and *UFGT*) increased after 3-6 days of cold storage, reconfirming previous data showing that the anthocyanin biosynthesis is regulated by the cold signaling pathway.

Some phenylpropanoids and anthocyanins are the major components of secondary metabolites that are affected by cold (Olsen *et al*., 2009; Li *et al*., 2015). Phenylpropanoids are diverse group of compounds derived from the carbon skeleton of the amino acid phenylalanine that is an end product of the shikimate pathway (Hermann and Weaver, 1999; Vogt, 2010; Fraser and Scheel, 2011). Phenylalanine is converted into 4-coumaroyl-CoA by three enzymes, i.e., phenylalanine ammonia-lyase (PAL), cinnamate 4-hydroxylase (C4H) and 4-coumarate:CoA ligase (4CL), in the general phenylpropanoid pathway (Supplementary Fig. S1). Chalcone synthase (CHS) catalyzes 4-coumaroyl-CoA to produce naringenin chalcone that is catalyzed into naringenin through a stereo specific isomerization reaction by chalcone isomerase (CHI). In next step, flavanone 3-hydroxylase (F3H) converts naringenin to dihydrokaempferol that can be catalyzed into dihydroquercetin by flavonoid 3’-hydroxylase (F3’H). Dihydroquercetin serves as a common precursor that can be catalyzed by either flavonol synthase (FLS) to form several flavonols or dihydroflavonol reductase (DFR) in the first step for anthocyanin biosynthesis (Winkel-Shirley, 2002). Anthocyanin synthase (ANS) then catalyzes the common step in the anthocyanin and proanthocyanidin (PA) biosynthetic pathway. Cyanidin is converted into cyanidin-O-glucoside and cyanidin-O-rutinoside by flavonoid 3-O-glucosyltransferase (3GT) and 3-O-rhamnosyltransferase (RT), respectively. Leucoanthocyanidin reductase (LAR) and anthocyanidin reductase (ANR) can produce proanthocyanidins (PAs) such as catechin and epicatechin (Abrahams *et al*., 2003; Xie *et al*., 2003; Matsui *et al*., 2016).

Tartary buckwheat (*Fagopyrum tataricum*) is an annual plant in the family Polygonaceae that has been considered to be an alternative crop or minor cereal in Asia, Europe, North America and South Africa. Recently, tartary buckwheat has become known as a rich source of numerous health-benefiting compounds such as vitamins B1, B2 and B6, rutin, quercetin, chlorogenic acid, anthocyanins and proanthocyanidins (Bonafaccia *et al*., 2003; Suzuki *et al*., 2005; Kim *et al*., 2007). In particular, rutin is the predominant flavonoid in tartary buckwheat that reaches 50-60 mg g^-1^ dry weight (Kim *et al*., 2006). Despite the nutritional value of tartary buckwheat, the transcriptome and metabolome responses to abiotic stress in this plant are still poorly understood. In this study, we identified the changes that occur in the metabolome and transcriptome of tartary buckwheat in response to cold, focusing on the flavonoid biosynthetic pathway. We studied the dynamic transcriptional levels of the phenylpropanoid biosynthetic genes to cold. By analyzing both the primary and secondary metabolites of tartary buckwheat without or with cold, we showed metabolic responses to environmental stresses.

## Materials and Methods

### Plant material and cold treatment

Tartary buckwheat cultivar “Hokkai T8” was provided by the Hokkaido Agricultural Research Center (Hokkaido, Japan). Dehulled tartary buckwheat seeds were sterilized with 70% ethanol for 30 s and 4% (v/v) bleach solution for 15 min, and then rinsed several times in sterile water. These seeds were grown on agar plates containing sucrose-free half Murashige and Skoog (1/2 MS) medium with 2.5 mM MES, pH 5.7, and 0.8% agar at 25°C under light/dark (16/8 h) conditions. Six-day-old light-grown plants were incubated at 4°C for varying periods with white fluorescent light for cold treatments. The samples were immediately frozen in liquid nitrogen and then stored at -80°C for RNA isolation or freeze-dried for high-performance liquid chromatography (HPLC) analysis.

### Illumina sequencing of the transcriptome

Total RNA was isolated from the frozen seedlings of tartary buckwheat using an RNeasy Mini Kit (Qiagen, Valencia, USA) and cleaned by ethanol precipitation. We removed the rRNAs in the total RNA using a ribo-zero rRNA removal kit (Epicentre, RZPL11016) and constructed a cDNA library for RNA sequencing using a TruSeq stranded total RNA sample prep kit-LT set A and B (Illumina, RS-122-2301 and 2302) according to the manufacturer’s instructions (Illumina, San Diego, CA, USA). The cDNA library was sequenced in 76 bp length paired-end (PE) reads in an Illumina NextSeq500 sequencer (Illumina Inc., San Diego, CA, USA) to produce 69,570,892 raw sequencing reads.

### De novo assembly and annotation of the tartary buckwheat transcriptome

The quality-trimmed reads of tartary buckwheat RNAs were assembled as contigs of the tartary buckwheat transcriptome using the Trinity software package (http://trinityrnaseq.github.io) (Haas *et al*., 2013). The characteristic properties including the N50, average, maximum, and minimum lengths of the assembled contigs were calculated using Transrate software (http://hibberdlab.com/transrate) (Smith-Unna *et al*., 2016). We clustered the tartary buckwheat transcriptome contigs based on sequence similarity using CD-HIT-EST software (http://weizhongli-lab.org/cd-hit) (Fu *et al*., 2012). To infer the biological functions of tartary buckwheat transcripts, we performed a homology search of the transcripts in a number of public protein and nucleotide databases. Transcript lists and sequences are presented in Additional File 1 and Additional File 2, respectively. The functional category distributions of tartary buckwheat transcripts in terms of Gene Ontology (GO) and COG were evaluated using the results of the homology search. COG functional category information attached to the COG proteins that fit the search parameters was used to determine the for determining COG functional category distribution, and GO information attached to the hit UniProt proteins was collected and re-analyzed using the WEGO tool (http://wego.genomics.org.cn) (Ye *et al*., 2006) in terms of the level for the three GO categories.

### Differentially expressed gene analysis

To quantify tartary buckwheat transcript expressions, we aligned preprocessed quality-trimmed reads on the tartary buckwheat transcript sequences and calculated the expression values with the aligned read counts for each transcript. Bowtie2 (http://bowtie-bio.sourceforge.net/bowtie2) software (Langmead and Salzberg, 2012) was used to align the quality-trimmed reads on the transcript sequences, and eXpress (http://bio.math.berkeley.edu/eXpress) software (Roberts and Pachter, 2013) was used to evaluate gene expression, in terms of fragments per kilobase of exon per million mapped fragments (FPKM) from the aligned results. The FPKM method provides a comparison between genes within a sample or between samples by normalizing the amount of sequencing for samples and gene length bias during gene expression evaluation.

### Real-time RT-PCR

Total RNA was isolated from frozen buckwheat seedlings using a total RNA mini kit (Geneaid Biotech Ltd., New Taipei City, Taiwan). Real-time RT-PCR was conducted using a QuantiTect SYBR Green RT-PCR kit (Qiagen, Valencia, USA) in a CFX96TM Real-Time PCR detection system (Bio-Rad, Hercules, CA). Gene expression was normalized to that of the histone H3 gene as a housekeeping gene. All real-time RT-PCR assays were conducted in triplicate biological replications and subjected to statistical analysis. Real-time RT-PCR conditions and primer sequences are provided in Supplementary Table S1.

### GC-TOFMS analysis of polar metabolites

Polar metabolites were extracted and silylated as described by Kim *et al* (2013). Briefly, metabolites were extracted with a mixed solvent of methanol: water: chloroform (2.5:1:1). Ribitol solution (60 μL, 0.2 mg/ml) was added as an internal standard (IS). The methanol: water phase containing the polar metabolites was dried in a centrifugal concentrator (CVE-2000, Eyela, Japan) and a freeze dryer. Methoxime and trimethylsilyl (TMS) etherification was performed for GC-TOFMS analysis. GC-TOFMS was performed using an Agilent 7890A gas chromatograph (Agilent, Atlanta, GA, USA) coupled to a Pegasus HT TOF mass spectrometer (LECO, St. Joseph, MI). The GC-MS conditions were as follows: column, 30-m **×** 0.25-mm I.D. fused-silica capillary column coated with 0.25-μm CP-SIL 8 CB low bleed (Varian Inc., Palo Alto, CA, USA); carrier gas, helium; gas flow rate, 1.0 mL/min; injector temperature, 230°C; transfer line temperature, 250°C; ion-source temperatures, 200°C; and detector voltage, 1700 V. The temperature program was as follows: initial temperature of 80°C for 2 min, followed by an increase to 320°C at 15°C/min, and a 10 min hold at 320°C.

### Extraction and HPLC analysis of anthocyanin

Anthocyanins were extracted from 100 mg dried powder with 2 mL water: formic acid (95:5 v/v) in a sonicator for 20 min, and the supernatant was filtered through a 0.45 μm hydrophilic PTFE syringe filter (Ø, 13 mm, Advantec, Tokyo, Japan) in a brown vial. Anthocyanins were detected at a wavelength of 520 nm using an Agilent 1200 series HPLC (Santa Clara, CA, USA) equipped with a Synergi 4 μm POLAR-RP 80A column (250 × 4.6 mm i.d., particle size 4 μm; Phenomenex, Torrance, CA, USA) and a Security Guard Cartridges Kit AQ C18 column (Phenomenex, Torrance, CA, USA). The mobile phase consisted of a mixture of (A) water: formic acid (95:5, v/v) and (B) acetonitrile: formic acid (95:5, v/v). The gradient program was set as follows: 5–10% solvent B from 0–8 min; 10–13% solvent B from 8–13 min; kept constant at 13% solvent B until 15 min; 13–15% solvent B from 15–18 min; kept constant at 15% solvent B until 25 min; 15–18% solvent B from 25–30 min, kept constant at 18% solvent B until 35 min; 18–21% solvent B from 35–40 min; kept constant at 21% solvent B until 45 min; quickly decreasing to 5% solvent B at 45.1 min; and kept constant at 5% solvent B until 50 min.

### Extraction and HPLC analysis of phenylpropanoids

For the phenylpropanoid analysis, 100 mg dried samples were extracted with 3 mL 80% methanol at 25°C for 1 h. After centrifugation at 12,000 × *g* for 10 min, the extracts were filtered through a 0.45 μm PTFE syringe filter (Advantec DISMIC-13HP, Toyo Roshi Kaisha, Ltd., Tokyo, Japan). Phenylpropanoids were separated using a Futecs model NS-4000 HPLC apparatus (Daejeon, Korea) with a C18 column (250 × 4.6 mm, 5 μm; RStech, Daejeon, Korea) and monitored at 280 nm. The mobile phase consisted of a mixture of 0.15% acetic acid (solvent A) and 100% methanol (solvent B). The initial mobile phase composition was as follows: 5% solvent B, followed by a linear gradient from 0 to 80% solvent B over 93 min and then a hold at 5% solvent B for an additional 5 min. The column was maintained at 30°C; the flow rate was 1.0 mL/min, and the injection volume was 20 μL. Different compounds were quantified based on peak areas, and the concentrations were calculated as equivalents of representative standard compounds.

### DPPH assay

The 2,2-diphenyl-1-picrylhydrazyl (DPPH) scavenging assay of buckwheat seedlings was conducted as described by Al-Dhabi *et al*. (2015). We prepared 0.15% DPPH in ice-cold methanol. The reaction mixture contained 4 mL methanol and various concentrations of extracts (20-100 *μ*L in a 1 mL volumn). Two hundred microliters DPPH solution was added followed by incubation at room temperature for 30 min in the dark. After incubation, the absorbance was measured at 517 nm. The DPPH radical scavenging activity was calculated by the following formula: DPPH radical scavenging activity (%) = [(*A*0 - *A*1)/*A*0 *\**100], where *A*0 is the absorbance of the control at 30 min and *A*1 is the absorbance of the sample at 30 min. All samples were analyzed in triplicate biological replications.

### Statistical analysis

Statistical analysis was performed with SPSS statistics 22 software using Student’s *t* test or an analysis of variance (ANOVA) with Tukey’s honestly significant difference test. All data are the mean values and standard deviation or standard error of triplicate experiments.

## Results

### Phenotype of tartary buckwheat plants in response to cold

To determine whether tartary buckwheat plants are affected by cold, we analyzed the morphology of *F. tataricum* ‘Hokkai T8’ seedling from 2 to 8 days in response to cold. *F. tataricum* ‘Hokkai T8’ seedlings have a light green hypocotyl and green cotyledons under normal growth conditions, while the seedlings exposed to the cold have a red hypocotyl and reddish green cotyledons with a color that was more intense at 8 day after exposure (DAE) to cold (Fig. 1A). The morphology of the tartary buckwheat seedlings was also greatly affected by cold (Fig. 1B and 1C). The fresh weight and length of seedlings grown at 25°C increased with time. The fresh weight of the seedlings increased more than 2-fold in the seedlings at 8 DAE compared with the seedlings at 0 DAE. However, there were no significant increases in the fresh weight and length of the seedlings grown under cold for 2, 4, 6 and 8 DAE.

**Fig. 1.**
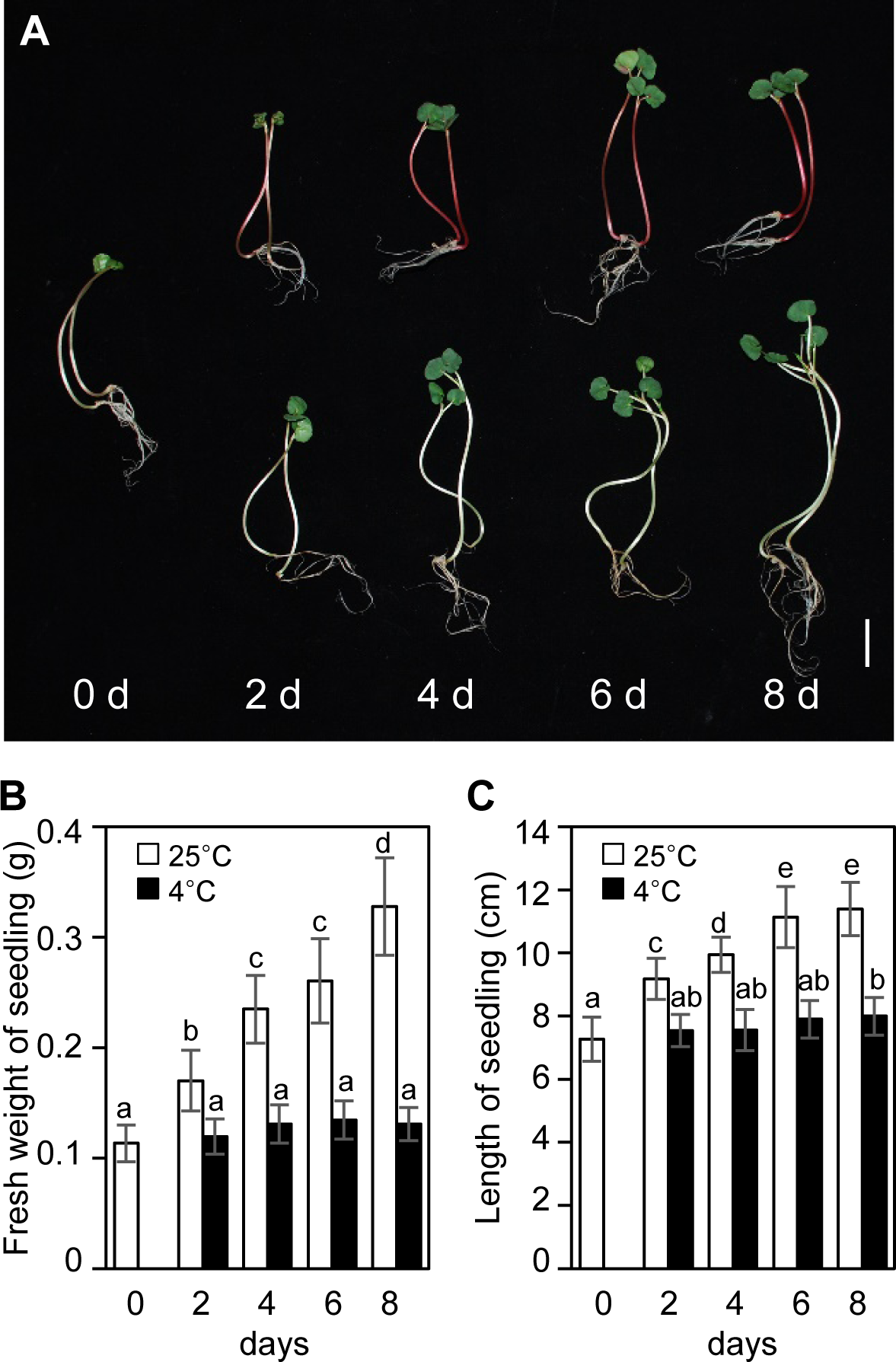
Phenotype of tartary buckwheat plants in response to cold. (A) Morphology of the tartary buckwheat seedlings in response to cold. Six-day-old light-grown plants were treated for 2, 4, 6, and 8 days at either 4°C (upper panel) or 25°C (bottom panel) before harvesting plant materials. Scale bars = 1 cm. (B) Fresh weight and (C) length of seedling from 0 to 10 days under 25°C or 4°C. Mean and SD values were determined from 20 seedlings. Bars with different letters indicate significant differences at P < 0.05 by one-way ANOVA with the Tukey’s honestly significant difference test.

### RNA-sequencing analysis of the cold transcriptome in tartary buckwheat plants

As shown in Fig. 1, cold stress could affect the morphology of the tartary buckwheat seedlings. We hypothesized that the cold could modulate the metabolite biosynthetic genes for the adaptation of buckwheat to cold stress. To identify the cold-regulated transcriptome in tartary buckwheat, we conducted RNA-sequencing analysis of tartary buckwheat with cold at 4°C for 0, 4, 24 and 48 h using an Illumina NextSeq500 sequencer. Transcriptome analysis of the tartary buckwheat seedlings identified 65,724 transcripts with a mean size of 737 bp and N50 of 1,137 bp (Supplementary Table S2). The transcripts were annotated based on the BLASTX algorithm against the non-redundant (NR) protein database and nucleotide (NT) database with an E-value cutoff of 1 x 10^-5^ (Supplementary Table S3). Of the total 65,724 transcripts, 48,868 transcripts were identified, representing approximately 74.35% of all the assembled transcripts. We extracted the genes whose cold-responsive expression were increased over 2-fold with a false discovery rate (FDR) < 0.05 (Table 1). We found that 3,132, 4,595, and 1,597 transcripts with FDR < 0.05 were differentially expressed genes in the cold treated-sample after for 4, 24 and 48 h compared with the untreated plants, respectively. Comparative genome-wide expression analysis showed that 406, 762 and 583 transcripts were increased more than 2-fold with FDR <0.05 under cold treatment for 4, 24 and 48 h compared with the untreated plants, respectively. RNA sequencing of cold-treated buckwheat seedlings showed that a total of 1,203 transcripts were upregulated in response to cold at a minimum of one time point during the course of the experiment. Among them, we selected 10 candidate transcripts encoding enzymes related to the phenylpropanoid biosynthetic pathway that were highly similar to those of *Arabidopsis thaliana* (Fig. 2A). Heatmap analysis of the these genes showed that the *FtPAL1, FtPAL2, Ft4CL3, FtATOMT1* and *FtHCT* transcripts accumulated at 4 h in response to cold, reached a peak level at 24 h and then declined. Alternatively, *FtCHS, FtCHIL, FtF3H, FtFLS1* and *FtANS* transcripts, which are downstream genes of the phenylpropanoid biosynthetic pathway, showed their maximal cold response at 48 h.

**Table 1:** Summary of differentially expressed genes under cold treatment at 4°C. FDR < 0.05 and fold change > 2.0.

|  | All transcripts | Up-regulated transcripts |
| --- | --- | --- |
| Tested transcripts | 59,565 | - |
| DEGs at 4 h | 3,132 | 406 |
| DEGs at 24 h | 4,595 | 762 |
| DEGs at 48 h | 1,597 | 583 |
| DEGs at least 1 condition | 5,415 | 1,203 |

**Fig. 2.**
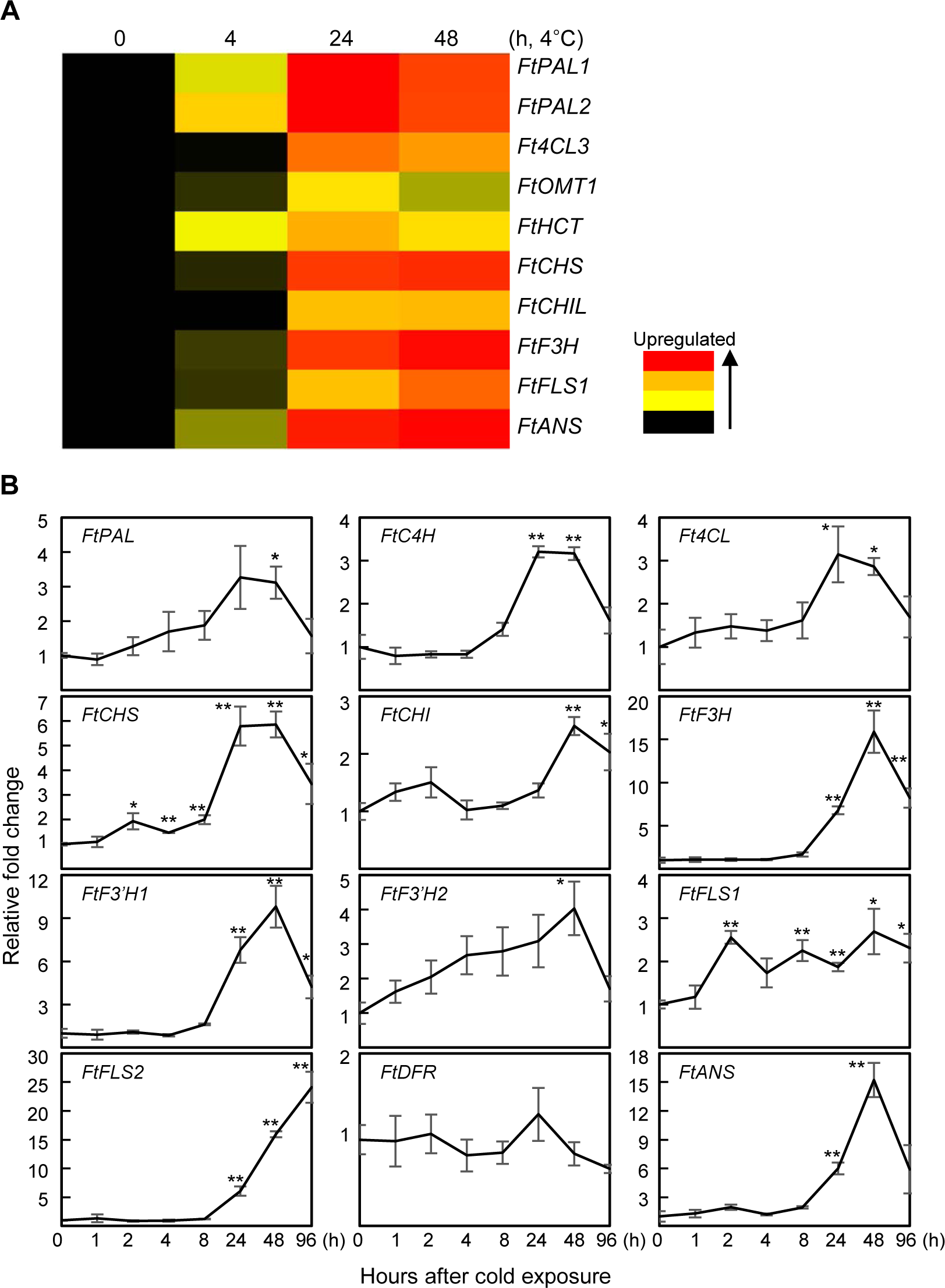
Expression of the phenylpropanoid biosynthetic genes in tartary buckwheat plants in response to cold. (A) Heatmap analysis of the phenylpropanoid biosynthetic genes in tartary buckwheat plants in response to cold. The response of the phenylpropanoid biosynthetic genes was obtained from RNA-sequencing analysis. The log_2_ fold change values for each time point were calculated. Scales indicate the color assigned to each log2 fold change. (B) QRT-PCR analysis of the phenylpropanoid biosynthetic genes in tartary buckwheat plants in response to cold. Six-day-old light-grown plants were treated at 4°C for varying periods of time (hour), and total RNA isolated was subjected to quantitative RT-PCR. Relative fold changes were plotted after normalization to histone H3. Mean values and SE from triplicate biological experiments are plotted. Asterisks indicate significant differences compared with the plant at 0 h using Student’s t test (*P < 0.05; **P < 0.01). PAL, phenylalanine ammonia-lyase; C4H, cinnamate 4-hydroxylase; 4CL, 4-coumarate-CoA ligase; OMT, O-methyltransferase; HCT, hydroxycinnamoyl-CoA shikimate/quinate hydroxycinnamoyl transferase; CHS, chalcone synthase; CHI, chalcone isomerase; CHIL, chalcone isomerase like; F3H, flavanone-3-hydroxylase; F3’H, flavonoid-3’-hydroxylase; FLS, flavonol synthase; DFR, dihydroflavonol reductase; ANS, anthocyanin synthase.

### Expression of phenylpropanoid biosynthetic genes in tartary buckwheat plants in response to cold

Our RNA sequencing data demonstrated that the expression of the phenylpropanoid biosynthetic genes was significantly induced by cold. These data are consistent with previous studies indicating that cold stress could affect the expression of metabolite biosynthetic genes in various plants such as *Arabidopsis* (Catala *et al*., 2011), blood orange (Crifo *et al*., 2012), purple kale (Zhang *et al*., 2012) and tobacco (Huang *et al*., 2012).

To identify whether the phenylpropanoid biosynthetic genes are affected by cold stress, we analyzed the expression of the phenylpropanoid biosynthetic genes in response to cold using quantitative real-time RT-PCR (Fig. 2B). The expression of most phenylpropanoid biosynthetic genes significantly increased in the cold treated buckwheat seedlings with the exception of *FtDFR*, which is expressed at a very low level in response to cold. *FtPAL, FtC4H* and *Ft4CL* exhibited maximal cold response at 24 h with induction levels more than 3.3-, 3.2- and 3.1-fold, respectively. *FtCHS, FtCHI, FtF3H, FtF3’H1, FtF3’H2, FtFLS1* and *FtANS* showed maximal cold response at 48 h with induction levels more than 5.9-, 2.5-, 15.9-, 9.8-, 4.0-, 2.7- and 15.2-fold, respectively. In particular, the expression of *FtANS*, the downstream gene of the phenylpropanoid biosynthesis pathway, was notably upregulated in response to cold. *FtFLS2*, which is a key gene of the flavonol biosynthetic pathway, was upregulated more than 24-fold after 96 h of cold treatment. These data are consistent with the cold transcriptome data of tartary buckwheat showing that cold induces the expression of several phenylpropanoid biosynthetic genes.

### Contents of metabolites in tartary buckwheat plants in response to cold

To identify the change of the primary metabolites in response to cold, we performed comprehensive metabolic profiling of the primary metabolites in tartary buckwheat grown during cold for 2, 5 and 8 days. A total of 44 metabolites including sugars, amino acids and organic acids were selected using direct comparison of the sample by GC-TOFMS analysis. The sugars and their derivatives significantly increased in cold-treated plants, with the exception of galactose, while most of the amino acids and their derivatives remain nearly unchanged or slightly decreased in the plants grown at 4°C compared with the plants grown at 25°C (Fig. 3). Xylose, mannitol, glucose, sucrose, inositol, mannose and fructose contents increased more than 8.7-, 2.7-, 3.3-, 2.6-, 12.4-, 6.3- and 5.8-fold after 8 days of cold treatment compared with the plants grown at 25°C, respectively (Fig. 3A). Threonic acid, a sugar acid derived from threose that is a four-carbon monosaccharide, also increased more than 2.0-fold after 8 days of cold treatment compared with the plants grown at 25°C. Among the sugars and their derivatives, the maltose content notably increased more than 20.6-fold in response to cold.

**Fig. 3.**
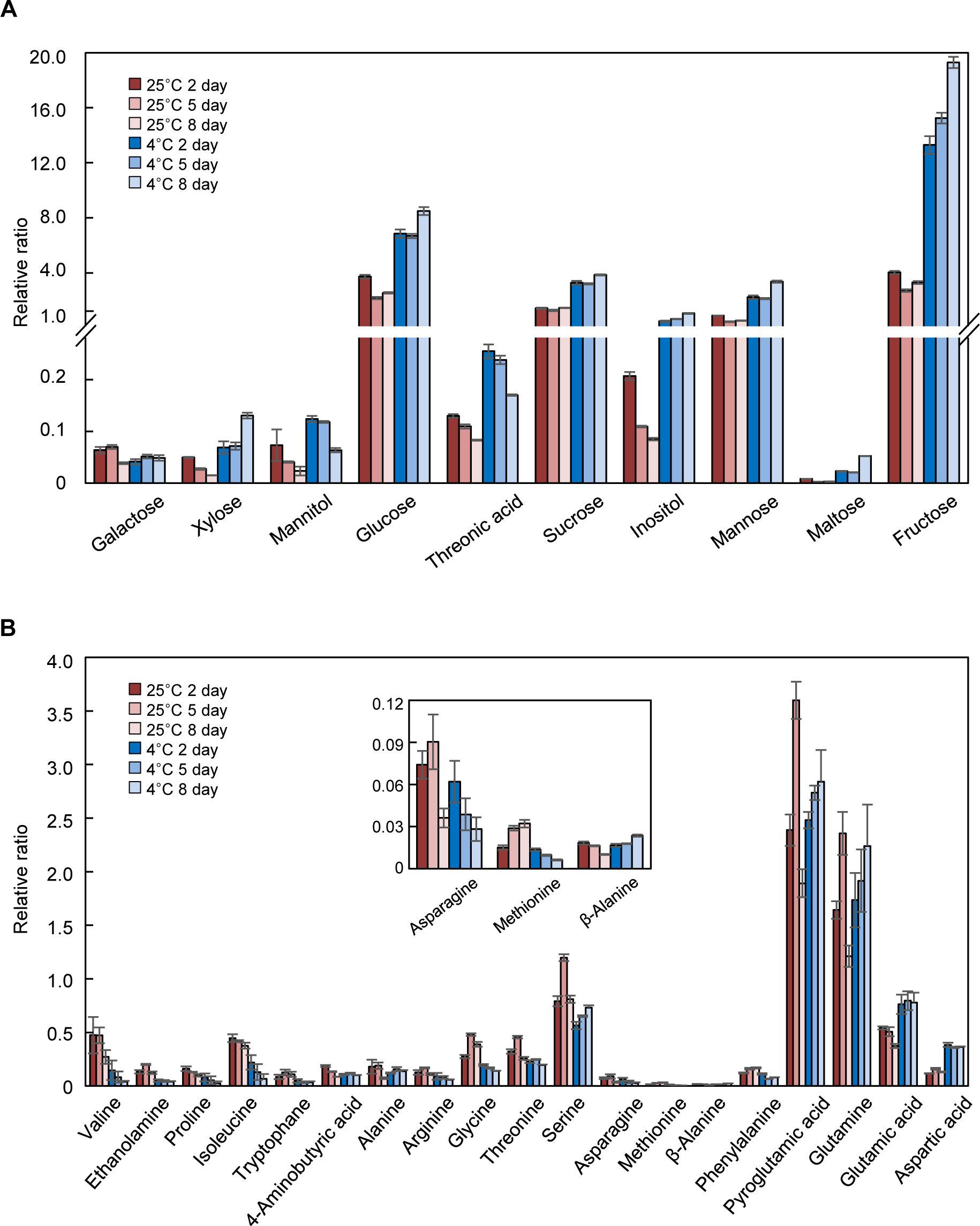
Contents of metabolites in tartary buckwheat plants in response to cold. (A) Sugars and their derivatives. (B) Amino acids and their derivatives. Six-day-old light-grown plants were treated at either 25°C (red) and 4°C (blue) for varying periods of time (day). Each value is the mean of triplicate biological experiments, and error bars indicate SD.

In primary metabolic profiling, 19 amino acids and their derivatives were detected in tartary buckwheat (Fig. 3B). Among the 19 amino acids and their derivatives, 11 of these compounds including valine, ethanolamine, proline, isoleucine, tryptophan, arginine, glycine, threonine, serine, asparagine, methionine and phenylalanine decreased more than 0.14-, 0.32-, 0.23-, 0.17-, 0.33-, 0.53-, 0.36-, 0.76-, 0.90-, 0.77-, 0.19- and 0.46-fold after 8 days of cold treatment compared with the plants grown at 25°C, respectively. Alternatively, the contents of *β*-alanine, pyroglutamic acid, glutamine, glutamic acid and aspartic acid increased more than 2.32-, 1.50-, 1.84-, 2.08- and 2.75-fold after 8 days of cold treatment compared with the plants grown at 25°C, respectively. The 4-aminobutyric acid (GABA) content decreased in plants grown at 25°C, while the slightly decreased 4-aminobutyric acid content was unchanged in cold conditions.

Fourteen organic acids were detected in the tartary buckwheat (Supplementary Fig. S2). Some organic acids derived from the tricarboxylic acid (TCA) cycle such as citric acid, fumaric acid, succinic acid and malic acid increased more than 2.23-, 2.47-, 1.36- and 3.57-fold after 8 days of cold treatment compared with the plants grown at 25°C, respectively. Of the shikimic pathway-related organic acids, vanillic acid decreased 0.78-fold, while shikimic acid, *p*-hydroxybenzoic acid, quinic acid and ferulic acid increased 1.29-, 1.25-, 9.07- and 1.54-fold after 8 days of cold treatment compared with the plants grown at 25°C, respectively.

In summary, our data showed that the contents of carbon metabolism-associated metabolites such as sugars and their derivatives increased significantly in cold-treated plants, while the contents of nitrogen metabolism-associated metabolites such as amino acids and their derivatives decreased in cold-treated plants, indicating that tartary buckwheat underwent major changes in carbon-nitrogen metabolism in response to cold.

### Contents of phenylpropanoids in tartary buckwheat plants in response to cold

In general, the production of secondary metabolites is associated with the pathways of primary metabolism. Tartary buckwheat has only two anthocyanins that include cyanidin 3-O-glucoside and cyanidin 3-O-rutinoside (Thwe *et al*., 2014). To investigate the contents of anthocyanins and proanthocyanidins that are regulated by cold, we analyzed the anthocyanins and proanthocyanidins in tartary buckwheat grown in the cold for 2, 4, 6 and 8 days (Fig. 4). The contents of cyanidin 3-O-glucoside and cyanidin 3-O-rutinoside were significantly increased by cold treatment (Fig. 4A). Cyanidin 3-O-glucoside was not detected in plants grown at 25°C, while the cyanidin 3-O-glucoside content of the cold-treated buckwheat increased more than 11.3-fold from 0.03 mg g^-1^ DW at 2 DAE to 0.34 mg g^-1^ DW at 8 DAE following cold treatment. The cyanidin 3-O-rutinoside content of plants grown at 25°C did not change significantly during plant development, while the cyanidin 3-O-rutinoside content of the cold-treated buckwheat increased more than 6.3-fold from 0.67 mg g^-1^ DW at 0 DAE to 4.24 mg g^-1^ DW at 8 DAE following cold treatment.

**Fig. 4.**
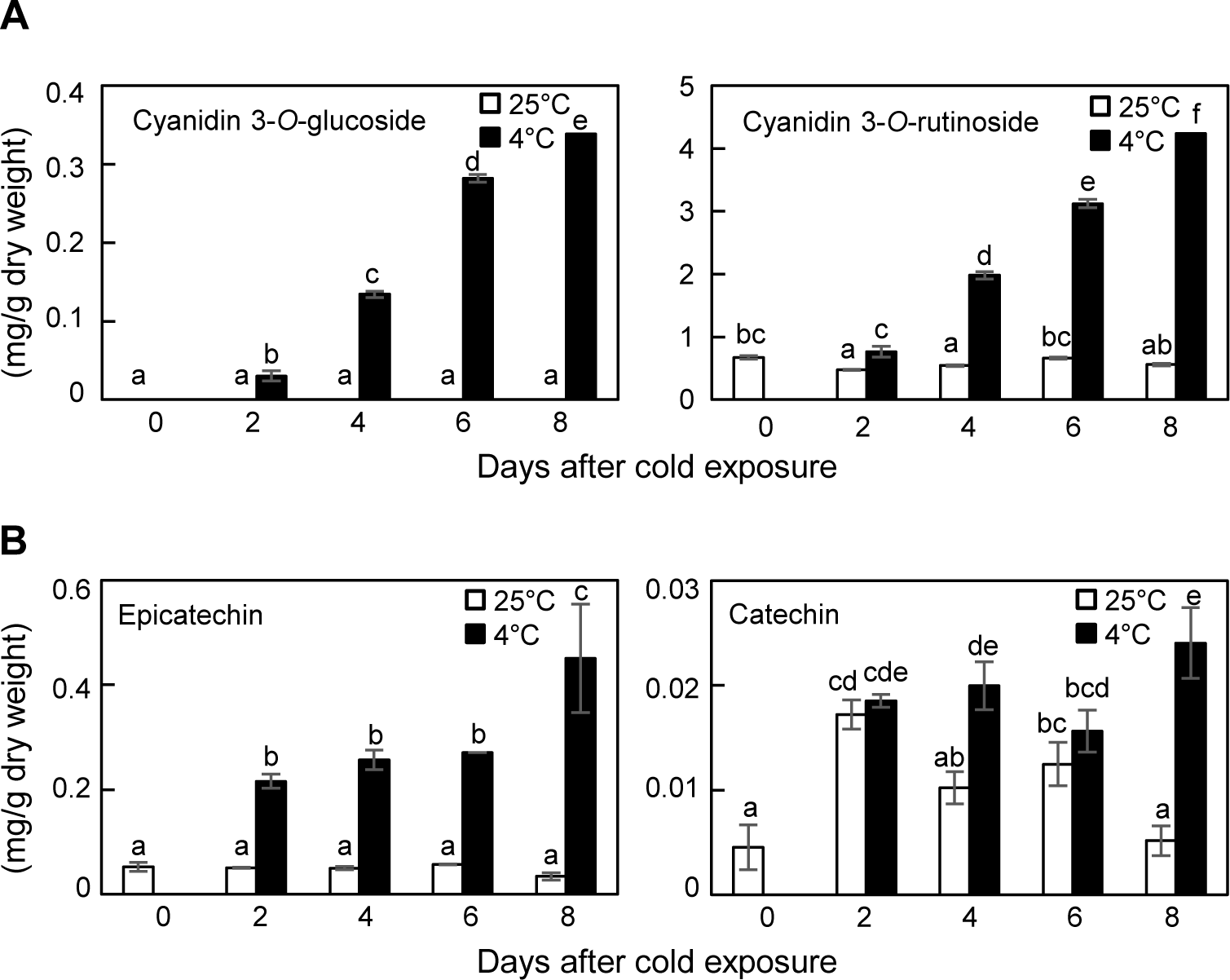
Accumulation of anthocyanins (A) and proanthocyanidins (B) in tartary buckwheat plants in response to cold. Six-day-old light-grown plants were treated at 4°C for varying periods of time (day). Each value is the mean of triplicate biological experiments, and error bars indicate SD. Bars with different letters indicate significant differences at P < 0.05 by one-way ANOVA with the Tukey’s honestly significant difference test.

Proanthocyanidin contents were also significantly increased in cold-treated plants (Fig. 4B). The epicatechin content of plants grown at 25°C did not change significantly during plant development, while the epicatechin content of the cold-treated buckwheat was increased more than 8.6-fold from 0.052 mg g^-1^ DW at 0 DAE to 0.451 mg g^-1^ DW at 8 DAE following cold treatment. In plants grown at 25°C, the catechin content exhibited its maximal level at day 2 and then declined. Alternatively, the catechin content of the cold-treated buckwheat increased more than 5.3-fold from 0.005 mg g^-1^ DW at 0 DAE to 0.024 mg g^-1^ DW at 8 DAE following cold treatment.

The chlorogenic acid content slightly increased more than 1.4-fold after 8 days of cold treatment compared with the plants grown at 25°C (Supplementary Fig. S3). The ferulic acid, rutin and quercetin contents did not differ significantly between plants grown at 25°C and those grown in the cold. Alternatively, the caffeic acid content was significantly reduced more than 4.3-fold after 8 days of cold treatment compared with the plants grown at 25°C.

### Radical scavenging activity of tartary buckwheat plants in response to cold

To investigate the effect of the radical scavenging activity by cold treatment, we performed the 2,2-diphenyl-1-picrylhydrazyl (DPPH) assay. The DPPH assay is a valid and easy method to evaluate the scavenging activity of flavonoids towards the stable radical DPPH (Seyoum *et al*., 2006). The radical scavenging activity of the cold-treated plants was significantly increased compared to that from the untreated-plants (Fig. 5). The radical scavenging activity of the untreated-plants did not change during the experimental period, while the radical scavenging activity of the cold treated-plants increased more than 5-fold compared with the untreated-plants. From this result, we predict that flavonoid accumulation by cold treatment leads to the enhancement of antioxidant activity.

**Fig. 5.**
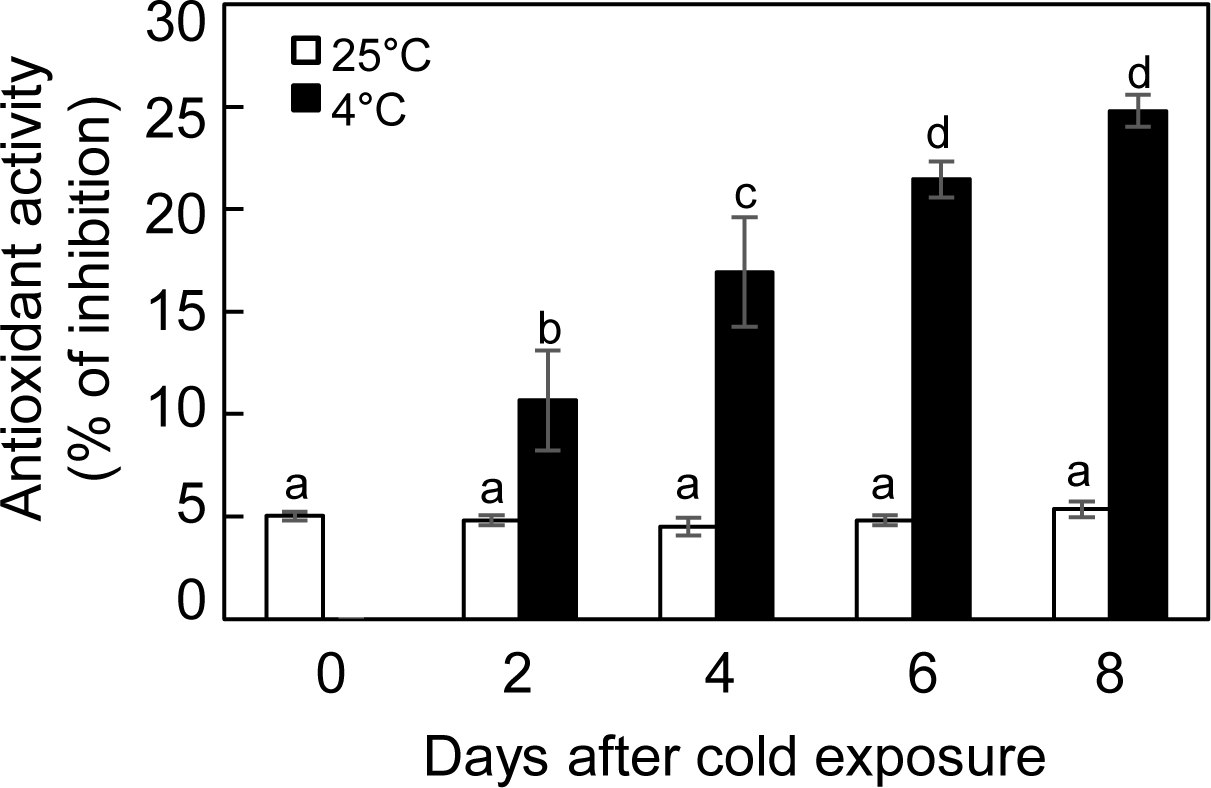
DPPH radical scavenging activity of tartary buckwheat plants in response to cold. Six-day-old light-grown plants were treated at 4°C for varying periods of time (day). Each value is the mean of triplicate biological experiments, and error bars indicate SD. Bars with different letters indicate significant differences at P < 0.05 by one-way ANOVA with the Tukey’s honestly significant difference test.

**Fig. 6.**
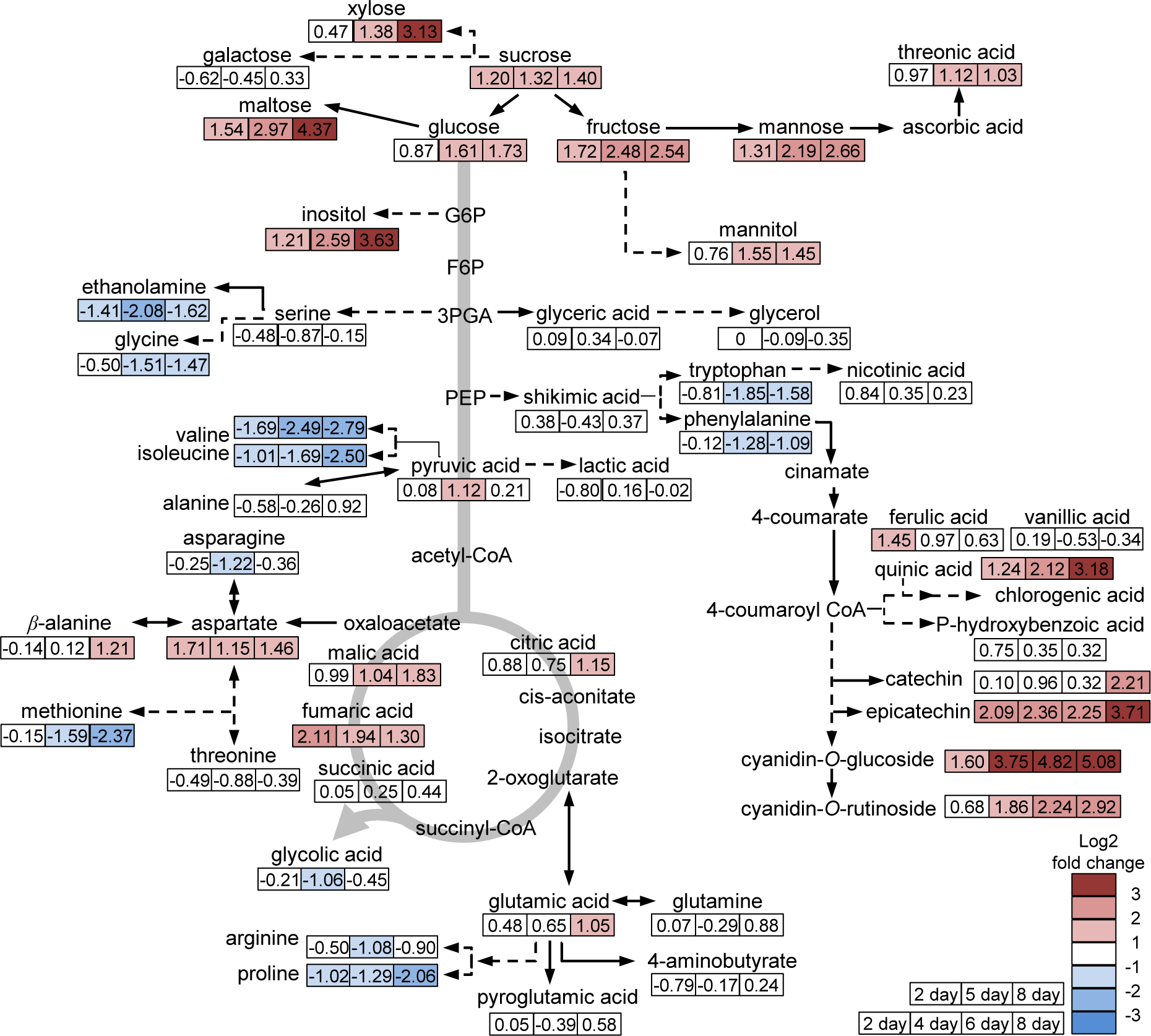
Overview of metabolic changes in response to cold in tartary buckwheat. The response ratio for each metabolite is given by cold-treated plants relative to cold-untreated plants in 2, 5, and 8 day or in 2, 4, 6, and 8 day.

## Discussion

During exposure to the cold, plants tend to biosynthesize primary and secondary metabolites to adapt to this condition. Certain sugars and amino acids, flavonoids and ascorbate accumulate to adjust homeostasis and scavenging reactive oxygen species (ROS). Anthocyanins are representative secondary metabolites that are affected by cold (Olsen *et al*., 2009; Li *et al*., 2015). In blood orange, the anthocyanin content of fruit exposed to low temperature sharply increased more than 8-fold after 6 days of cold treatment compared with the time zero plants (Crifo *et al*., 2012). The expression of anthocyanin biosynthetic genes was significantly upregulated after 3-6 days of cold storage. Real-time RT-PCR analysis revealed that the transcripts encoding putative NAC domains increased during cold storage from 3 days of storage until the end of the period under investigation, showing a greater than 20-fold change. The anthocyanin accumulation in the purple kale is strongly induced by low temperatures (Zhang *et al*., 2012). The expression of the anthocyanin biosynthetic genes *C4H, F3H, DFR, ANS* and *UFGT* were all enhanced under low temperatures. The R2R3 MYB transcription factor entitled *BoPAP1* and the bHLH transcription factor entitled *BoTT8* were extremely induced by low temperature. In this study, we identified that the anthocyanin content induced by cold correlates with the regulation of gene expression. The contents of cyanidin 3-O-glucoside and cyanidin 3-O-rutinoside, types of anthocyanins found in tartary buckwheat, were significantly increased by cold treatment (Fig. 4A). RNA-sequencing analysis and quantitative real-time RT-PCR data of tartary buckwheat showed that most of the phenylpropanoid biosynthetic genes including *FtPAL, FtC4H, Ft4CL, FtCHS, FtCHI, FtF3H, FtF3’H1, FtF3’H2, FtFLS1, FtFLS2* and *FtANS* were upregulated in response to cold (Fig. 2). However, the expression of *FtDFR*, the late anthocyanin biosynthetic gene, was not affected by cold in tartary buckwheat. These data differs from the upregulation of *DFR* by cold in maize (Christie *et al*., 1994), *Arabidopsis* (Zhang *et al*., 2011), blood orange (Crifo *et al*., 2012) and purple kale (Zhang *et al*., 2012). These results suggest that the other late anthocyanin biosynthetic genes could play major roles in anthocyanin biosynthetic pathway under cold conditions.

Of the metabolites that changed in response to cold, most of the sugars and their derivatives significantly increased more than 2.0-fold after 8 days of cold treatment compared with the plants grown at 25°C (Fig. 3A). The increases in the levels of sugars correlated with those in the anthocyanin biosynthetic pathway (Solfanelli *et al*., 2006). Sucrose affects the genes encoding for enzymes involved in the anthocyanin biosynthetic pathway in *Arabidopsis*. In particular, sucrose is an effective stimulator for the mRNA accumulation of late anthocyanin biosynthetic genes such as *DFR, ANS* and *3GT*. The *Arabidopsis pho3* mutant has a defect in the *SUCROSE-PROTON SYMPORTER 2* (*SUC2*) gene restricting sucrose transport. The mutant has a dwarf phenotype with growth retardation, leading to accumulation of soluble sugars, starch and anthocyanin. The transcriptome data of the *pho3* mutant showed an enhanced expression of genes encoding for the anthocyanin biosynthetic enzymes, as well as the transcription factors for the anthocyanin biosynthetic pathway, suggesting that the sugars are *in vivo* stimulators of anthocyanin biosynthesis (Lloyd and Zakhleniuk, 2004). Based on previously published data and our transcriptomic and metabolic profiling, we suggest that cold plays a role in the modulation of sugar synthesis, increasing the contents of sugars. The increased sugars might be act as signaling molecules to induce the expression of genes encoding anthocyanin biosynthetic enzymes, thus enhancing the accumulation of anthocyanins. However, we cannot rule out the possibility that cold directly modulate the anthocyanin biosynthetic pathway without sugar signaling.

## Supplementary data

Supplementary data are available at *JXB* online.

Table S1: Primers used in this work.

Table S2: Summary of the transcriptome of cold-treated tartary buckwheat.

Table S3: Summary of annotations of tartary buckwheat transcripts.

Fig. S1: Schematic representation of the phenylpropanoid biosynthetic pathway in tartary buckwheat plants.

Fig. S2: Contents of organic acids in response to cold in tartary buckwheat plants.

Fig. S3: Accumulation of phenolic compound in response to cold in tartary buckwheat plants.

## Acknowledgements

This research was supported by Basic Science Research Program through the National Research Foundation of Korea (NRF) funded by the Ministry of Education (2016R1A6A3A11935897). The authors declare that they have no competing interests.

## Data availability

Our Illumina RNA sequencing data for *Fagopyrum tataricum* have been deposited in the NCBI Short Read Archive database under accession number SRP132029.

